# Vascular and Lymphatic Dysregulation via Non-EndoMT *Col2a1* Signaling in Bisphosphonate-Related Osteonecrosis of the Jaw

**DOI:** 10.1101/2025.09.27.678999

**Authors:** Yixin Shi, Haiyang Sun, Zijian Guo, Xian Liu, Xibo Pei, Jian Wang, Fanyuan Yu, Xingchen Peng, Quan Yuan, Anjali P. Kusumbe, Junyu Chen

## Abstract

Bisphosphonate-related osteonecrosis of the jaw (BRONJ) is one of the most severe complications of antiresorptive therapy. Although impaired angiogenesis is commonly observed in BRONJ, the underlying vascular mechanisms remain poorly understood, impeding the development of effective treatments. In this study, we integrated whole-organ tissue-clearing imaging, single-cell transcriptomics, and proteomics to construct a multimodal atlas of the mandibular microenvironment in both mouse and human BRONJ. Our findings revealed gross vascular and lymphatic rarefaction in BRONJ, along with 4 endothelial subtypes uniformly exhibiting aberrant upregulation of *Col2a1*. Moreover, fibroblasts and macrophages emerged as key endothelial interactors, collectively underscoring dysregulation of the endothelium-matrix-immune axis. Unlike homeostatic *Dmp1*⁺/*Tfap2a*⁺ fibroblasts and *Stab1*⁺ macrophages, BRONJ lesions featured pathological *Lrrc15*⁺/*Chad*⁺ fibroblasts and *Il6*⁺ macrophages, which promote ectopic chondrogenesis and inflammation. Mechanistically, BRONJ activated the COL2A1-CD44 axis (EC-to-fibroblast/macrophage signaling) and COL2A1-SDC4 axis (EC-to-fibroblast signaling), whereas lineage tracing excluded endothelial-to-mesenchymal transition (EndoMT), implicating extracellular matrix remodeling as the driver of dysfunction. Cross-species validation in human mandibles further confirmed vascular-lymphatic dysregulation, inflammatory activation, and aberrant chondrogenesis. Overall, this study establishes vascular and lymphatic dysfunction as a central pathological hallmark of BRONJ, driving matrix remodeling dysregulation and immune-inflammatory imbalance, and identifies the COL2A1-CD44/SDC4 axis as a potential therapeutic target for BRONJ.

## Introduction

Bisphosphonate-related osteonecrosis of the jaw (BRONJ) is a serious complication of bisphosphonates, which are widely used to treat osteoporosis and cancer-related bone metastases (*1*). Clinically, BRONJ presents as non-healing bone exposure and necrosis with pain, infection, and soft-tissue involvement, often progressing to pathologic fractures and long-term jaw dysfunction (*2, 3*). Effective therapies for BRONJ remain lacking to date. Current management includes antiseptic mouth rinses, systemic antibiotics, resection of necrotic bone, and soft-tissue closure, yet recurrence remains high, largely owing to irreversible failure of vasculature repair (*4*). Although impaired angiogenesis has been implicated in BRONJ pathogenesis (*5*), the molecular basis of endothelial dysfunction and how it perturbs the local microenvironment remain undefined, representing a central knowledge gap.

Recent studies identify vascular endothelial dysfunction as central drivers of BRONJ progression (*6, 7*). Bisphosphonates hinder angiogenic repair by suppressing endothelial proliferatio and migration, and by reducing the regenerative capacity of endothelial progenitor cells (*8*). However, the heterogeneity of drug-exposed endothelial cell (EC) subtypes remains poorly defined. For instance, Emcn^hi^ CD31^hi^ type H vessels, represent a key subtype mediating the coupling of angiogenesis and osteogenesis (*9*); moreover, skeletal lymphatic vessels, as recently identified in our work (*10*), have been shown to play a pivotal role in osteogenesis under injury stress. However, most evidence relies on two-dimensional sections (*6, 11*), which lack spatial continuity and organ-level context, precluding a comprehensive view of the vasculature network. Recent study indicates bisphosphonates also impair lymphatic endothelium, suppress lymphangiogenesis and amplify immune-mediated inflammation (*7, 12*). Nonetheless, in situ characterization of bone lymphatics remains inadequate, complicating rigorous study and suggesting that non-tubular LYVE1 positivity may reflect macrophage rather than bona fide lymphatic vessels (*13*). Although endothelial injury is widely linked to the recalcitrant course of BRONJ, subtype-specific molecular programs, intercellular crosstalk, and downstream signaling networks remain unresolved. Moreover, despite extensive use of mouse and rat models (*14, 15*), human data are scarce and largely case-based (*16*), limiting cross-species generalization and mechanistic inference.

By integrating whole-organ 3D imaging, single-cell transcriptomics, and proteomics, based on murine and human disease samples, we constructed a multimodal atlas of the mandibular microenvironment in BRONJ. We delineate the molecular programs of endothelial subtypes and uncover their pathological crosstalk with fibroblasts and immune cells. We further identify the COL2A1-CD44 axis (EC-to-fibroblast/macrophage signaling) and COL2A1-SDC4 (EC-to-fibroblast signaling) axis as a central driver of endothelial dysfunction and define vascular-lymphatic dysregulation as a key pathogenic mechanism, thereby highlighting potential therapeutic avenues for BRONJ.

## Results

### Tooth-jawbone tissue-clearing imaging reveals vascular-lymphatic rarefaction in BRONJ

In recent years, advances in tissue-clearing imaging have greatly accelerated organ-level studies of the vasculature. For highly mineralized hard tissues such as bone and teeth, methods including TESOS (*17*), PEGASOS (*18*), and B-CLARITY (*19*) preserve endogenous fluorescence stability while providing fine microstructural detail and imaging clarity, yet show limited compatibility with multiplex exogenous immunolabeling (*20*). By enhancing antibody penetration into the central bone marrow region and applying hard-tissue clearing, our recent study demonstrated for the first time the existence of previously debated skeletal lymphatic vessels (*10*). Compared with long bones, the jawbone is an irregular bone with structure comprising both teeth and bone, which poses greater challenges for clearing (*21*). To address this, we established a clearing and imaging protocol optimized for the intact tooth-jawbone structure, incorporating collagenase digestion, graded isopropanol dehydration, and ethyl cinnamate clearing (Fig. 1A), which rendered the jaw optically transparent while allowing exogenous immunolabeling. Light-sheet microscopy captures the tooth-jawbone endothelium at the organ level, and confocal imaging of cleared sections resolves cellular and tissue details (Fig. 1A), together providing a multiscale view from single-cell to whole-organ.

**Fig. 1.**
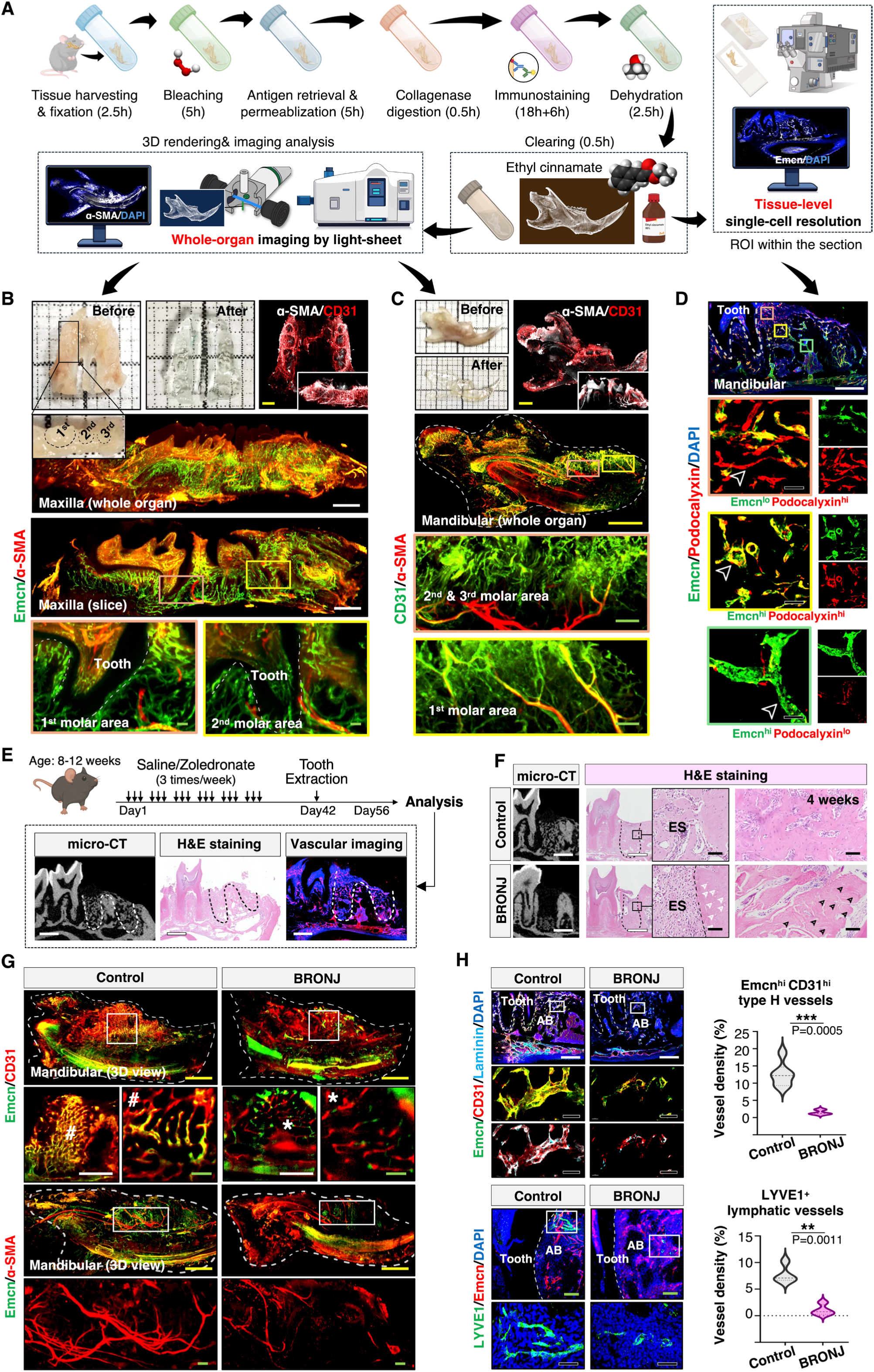
Tooth-jawbone tissue-clearing imaging reveals vascular and lymphatic alterations in BRONJ. (**A**) Schematic workflow of the tooth-jawbone clearing protocol for whole-organ imaging and region of interest (ROI) 3D imaging at single-cell resolution. (**B**) Maxillary bone before and after clearing, highlighting vasculature in the alveolar bone with high-resolution 3D reconstructions of the first and second molar regions; labeled with α-SMA, CD31, and Emcn. (**C**) Mandibular bone before and after clearing, showing high-resolution vascular structures of the first to third molar regions; labeled with α-SMA and CD31. (**D**) Imaging of vascular subtypes in the alveolar bone ROI, with Emcn, Podocalyxin labeling and DAPI nuclear staining; arrows mark representative subtypes. (**E**) Workflow for establishing the murine BRONJ model for subsequent analyses, including micro-CT, H&E staining, and vascular imaging of the extraction socket. (**F**) Micro-CT and hematoxylin and eosin (H&E) images of extraction sockets (ES) in control and BRONJ mandibles; arrows indicate empty bone lacunae. (**G**) Light-sheet imaging of cleared mandibles immunolabeled with Emcn, CD31, and α-SMA. (**H**) Immunofluorescence staining and semi-quantitative analysis of Emcn^hi^ CD31^hi^ type H vessels (*n=6*) and LYVE1⁺ lymphatic vessels (n=4) in control and BRONJ groups. **Scale bars:** yellow, 1000 µm; white, 500 µm; green, 100 µm; and black, 50 µm. ^ns^P >0.05; *P < 0.05; **P < 0.01; and ***P < 0.001. n represents biological replicates.

Using this approach, we visualized the whole-organ endothelium networks of both maxilla and mandible, and α-SMA⁺ perivascular cells (Fig. 1, B and C). Focusing on the molar region, we obtained 3D reconstructions of the 1^st^, 2^nd^, and 3^rd^ molars together with the surrounding dentoalveolar endothelium (Fig. 1, B and C). Furthermore, abundant neovessel clusters co-labeled by α-SMA, Emcn, and CD31 were observed in the extraction socket 2 weeks after first molar removal (fig. S1A). Higher-resolution imaging further revealed endothelial heterogeneity within the mandibular alveolar bone, distinguishing 3 vascular subtypes with distinct spatial distributions, including Emcn^lo^ Podocalyxin^hi^, Emcn^hi^ Podocalyxin^hi^, and Emcn^hi^ Podocalyxin^lo^ subsets (Fig. 1D). Thus, this workflow enables comprehensive endothelial mapping of the tooth-jawbone complex, providing a powerful tool for studying vascular and lymphatic pathology in jawbone diseases.

To investigate the endothelial contribution to BRONJ, we established a murine model by intraperitoneal injection of zoledronate, a first-line anti-osteoporotic drug(*22*), followed by bilateral extraction of the first mandibular molars (Fig. 1E, and fig. S1B). Zoledronate-treated mice exhibited reduced intraoperative bleeding, indicating vascular injury (fig. S1C), followed by soft-tissue ulceration and impaired wound healing after extraction (fig. S1D). Micro-CT and histology further confirmed the presence of necrotic bone in the BRONJ group (Fig. 1F, and fig. S1E). Whole-organ tissue-clearing imaging revealed a marked reduction of Emcn^hi^ CD31^hi^ type H vessels in BRONJ mice, as well as α-SMA^+^ mural cell-coverd arteries (Fig. 1G, and fig. S1F). At the tissue and single-cell level, both type H blood vessels and LYVE1⁺ lymphatic vessels showed pronounced rarefaction (Fig. 1H), accompanied by substantial loss of basement membrane components (fig. S1G). Together, these findings revealed profound vascular and lymphatic attenuation in BRONJ, motivating further exploration of the endothelial system as a key pathogenic mechanism and potential therapeutic target.

### A multidimensional endothelial atlas of cellular loss, signaling disruption, and metabolic imbalance in BRONJ

Single-cell transcriptomic profiling of healthy and BRONJ mouse mandibles yielded 34,350 high-quality cells (fig. S2, A to C), enabling construction of a comprehensive atlas spanning 12 major populations, including T cells, B and pre-B cells, natural killer (NK) cells, neutrophils, mast cells, macrophages, hematopoietic stem cells (HSCs), fibroblasts, erythrocytes, endothelial cells (ECs), and dendritic cells (DCs) (Fig. 2, A and B, and fig. S3A), with functional annotation validated by GO enrichment (fig. S3B). In BRONJ, ECs were markedly reduced (Fig. 2, C and D), accompanied by impaired vascular signaling (Fig. 2E). Among them, E-selectin plays a central role in vascular homeostasis (*23*) and stem cell niche regulation (*24*), and the *Sele* axis was nearly completely lost in BRONJ mice (Fig. 2E), further confirmed by immunolabeling (Fig. 2F). Protein-protein interaction (PPI) network and gene set enrichment analysis (GSEA) further revealed robust activation of osteogenic and inflammatory pathways in BRONJ tissues (Fig. 2G, and fig. S4A). Strikingly, under conditions of both numerical loss and signaling disruption, ECs uniquely exhibited concomitant upregulation of osteogenic and inflammatory transcriptional programs (Fig. 2H), whereas other cell types only displayed single-axis bias (fig. S4B). These findings highlight ECs as central hubs in BRONJ, simultaneously engaging osteogenic signaling and amplifying inflammatory injury.

**Fig. 2.**
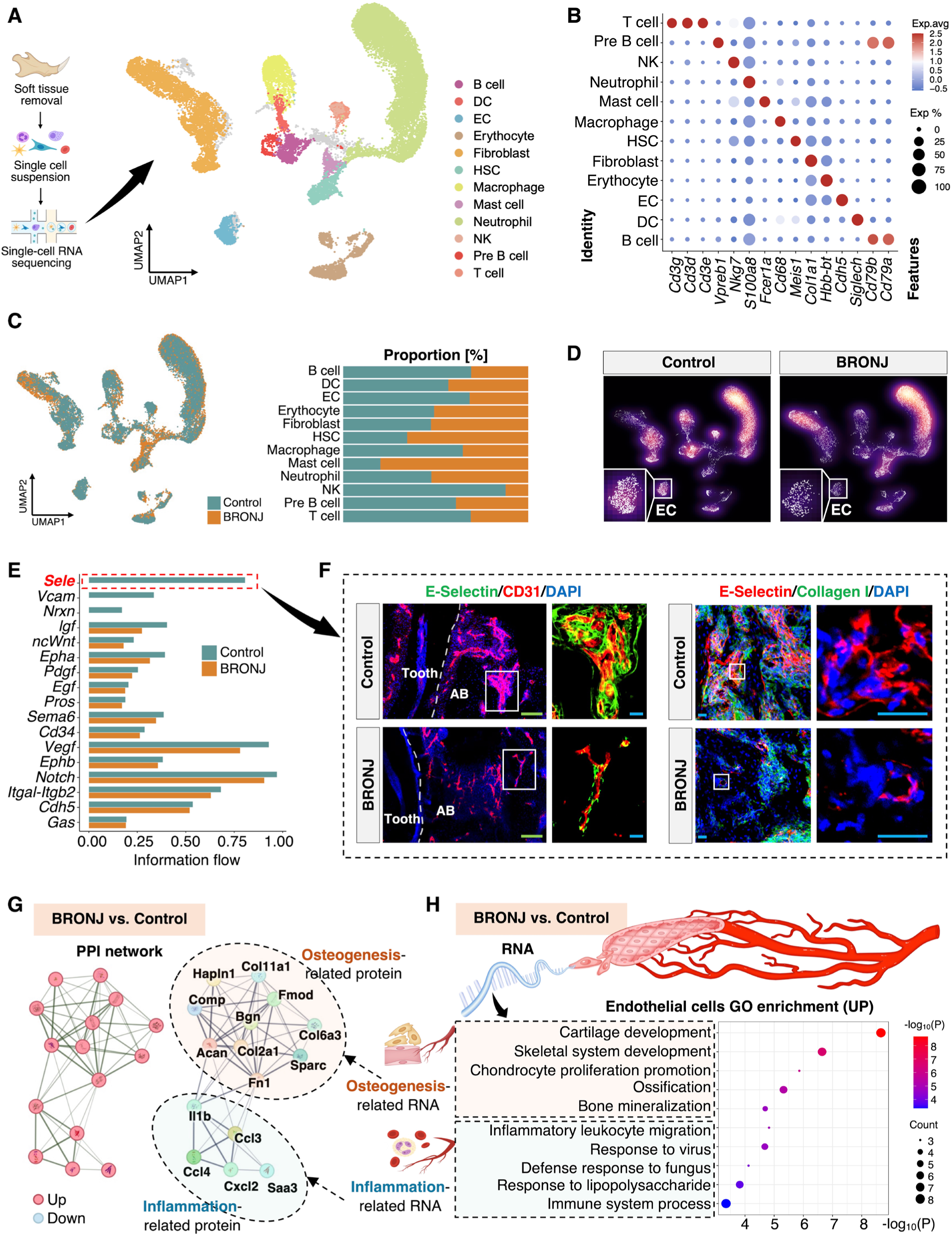
Single-cell atlas of healthy and BRONJ mandibles. (**A**) Experimental workflow and integrated UMAP of mandibular single-cell RNA sequencing from control (17,297 cells) and BRONJ (17,053 cells) groups. (**B**) Expression patterns of representative marker genes across 12 cell types. (**C**) Proportional distribution of cell types in control and BRONJ groups. (**D**) Cell density plots comparing groups, highlighting reduced EC density in BRONJ. (**E**) Information flow of endothelial signaling pathways in control and BRONJ. (**F**) Immunofluorescence imaging of control and BRONJ extraction sockets stained for E-Selectin, CD31, and Collagen I, with nuclear counterstaining by DAPI. (**G**) Protein-protein interaction (PPI) network changes highlighting osteogenesis- and inflammation-related proteins. (**H**) GO enrichment analysis of endothelial cell (EC) genes upregulated in BRONJ. **Scale bars**: green, 100 µm; and blue, 20 µm.

Recent advances in single-cell transcriptomics have greatly deepened our understanding of EC molecular programs and pathological networks, highlighting the remarkable heterogeneity of structure, function, and spatial distribution (*25, 26*). Building on this, we delineated EC heterogeneity in healthy and BRONJ mandibles and identified 4 major subtypes (Fig. 3A): *Gja4*⁺ arterial ECs (AECs), *Sele*⁺ venous ECs (VECs), *Fabp4*⁺ sinusoidal ECs (SECs), and *Lyve1*⁺ lymphatic ECs (LECs) (Fig. 3B). At single-cell resolution, we characterized these 4 endothelial subtypes (Fig. 3C); for the controversial LECs (*27*), we further validated their presence within jawbone and excluded macrophage contamination (fig. S5A).

**Fig. 3.**
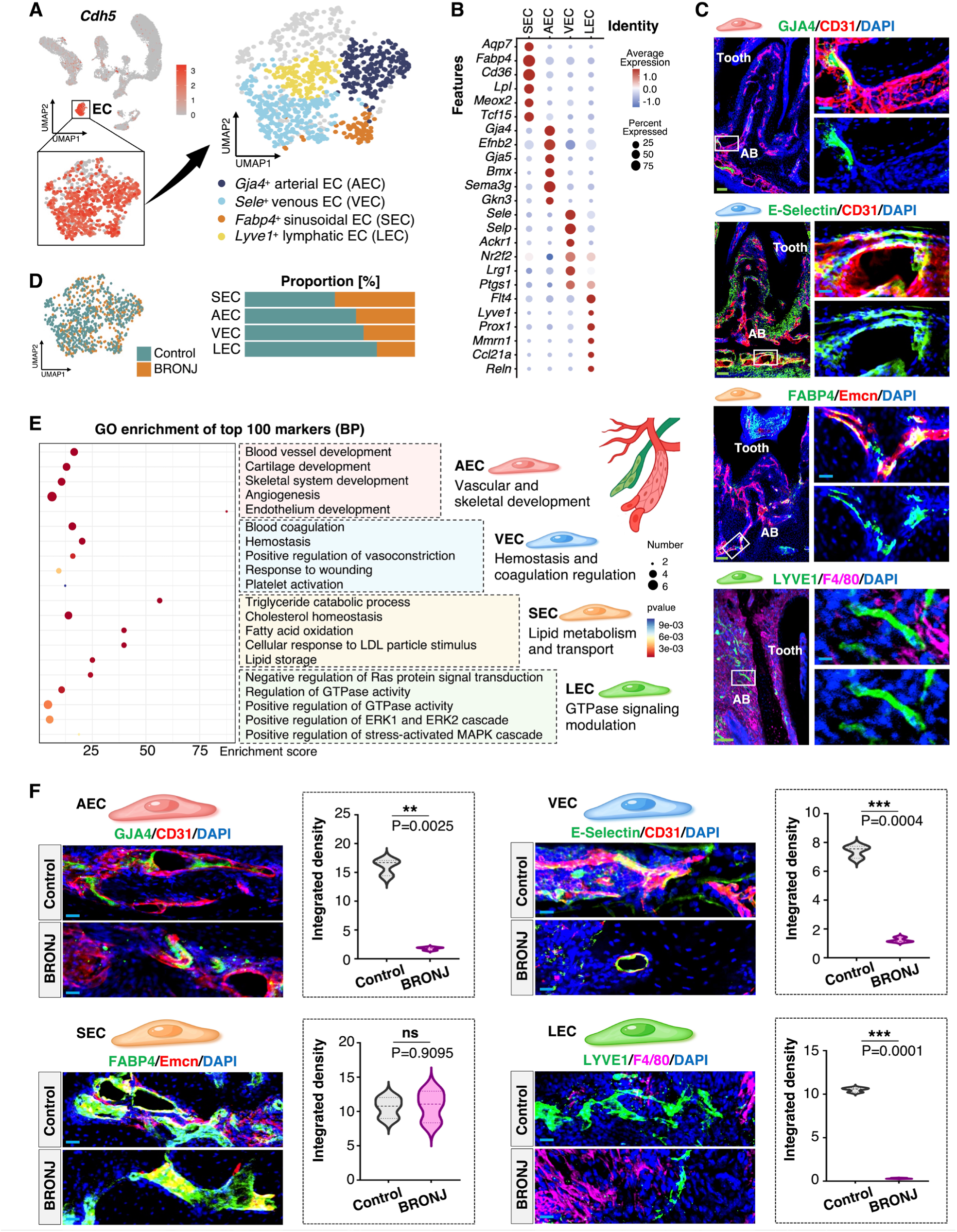
Single-cell atlas of endothelial heterogeneity and pathological alterations in BRONJ. (**A**) UMAP of *Cdh5*-marked ECs. (**B**) Expression of marker genes for EC subtypes. (**C**) Imaging of endothelial cell subtypes at single-cell resolution; AECs stained with GJA4 and CD31, VECs with E-Selectin and CD31, SECs with FABP4 and Emcn, and LECs with LYVE1 and F4/80; with nuclei counterstained by DAPI. (**D**) Proportional changes of EC subtypes between groups. (**E**) GO enrichment of the top 100 marker genes for each EC subtype. (**F**) Imaging and semi-quantitative analysis of EC subtype changes in control and BRONJ groups *(n=3)*; AECs, VECs, and SECs labeled as in **C**. **Scale bars**: green, 100 µm; and blue, 20 µm. ^ns^P >0.05; *P < 0.05; **P < 0.01; and ***P < 0.001. n represents biological replicates.

All 4 subtypes of EC showed reductions in BRONJ to varying degrees (Fig. 3D), indicating subtype-selective vulnerability. Functional annotation underscored distinct roles: AECs support vascular-bone development, VECs regulate hemostasis, SECs mediate lipid metabolism, and LECs maintain GTPase signaling (Fig. 3E). Trajectory analysis confirmed that these subtypes represent stable terminal states rather than transitional intermediates (fig. S5B). CellChat mapping further revealed markedly reduced vascular signaling in AECs, VECs, and LECs in BRONJ, whereas SECs partially preserved pathway activity (fig. S5C). GSEA results showed downregulation of angiogenic programs in most subtypes in BRONJ, while SECs retained relatively resistance to zoledronate-mediated angiogenic suppression (fig. S5D). Moreover, immunolabeling further validated significant depletion of all subtypes except SECs (Fig. 3F). ScMetabolism profiling revealed extensive EC reprogramming in BRONJ, spanning lipid, carbohydrate, and glycosaminoglycan metabolism (fig. S5E), all pathways closely associated with endothelial barrier integrity (*28*). Together, these findings define a multidimensional pathological network in BRONJ, in which endothelial loss, signaling disruption, and metabolic imbalance converge to drive disease progression.

### EC-stromal remodeling and aberrant chondrogenic fibroblasts in BRONJ

After establishing the central role of ECs in BRONJ, we next focused on their downstream stromal partners. CellChat analysis revealed globally enhanced intercellular communication in the BRONJ microenvironment, with fibroblasts emerging as the primary recipients of endothelial signaling (Fig. 4, A and B). This heightened communication was accompanied by coordinated transcriptional changes, as both ECs and fibroblasts co-upregulated a set of genes including *Col2a1*, *Comp*, *S100a9*, and *S100a8* (fig. S6A). Further, fibroblast-upregulated genes were enriched in cartilage development pathways (fig. S6B), indicating a shift toward chondrogenic remodeling.

**Fig. 4.**
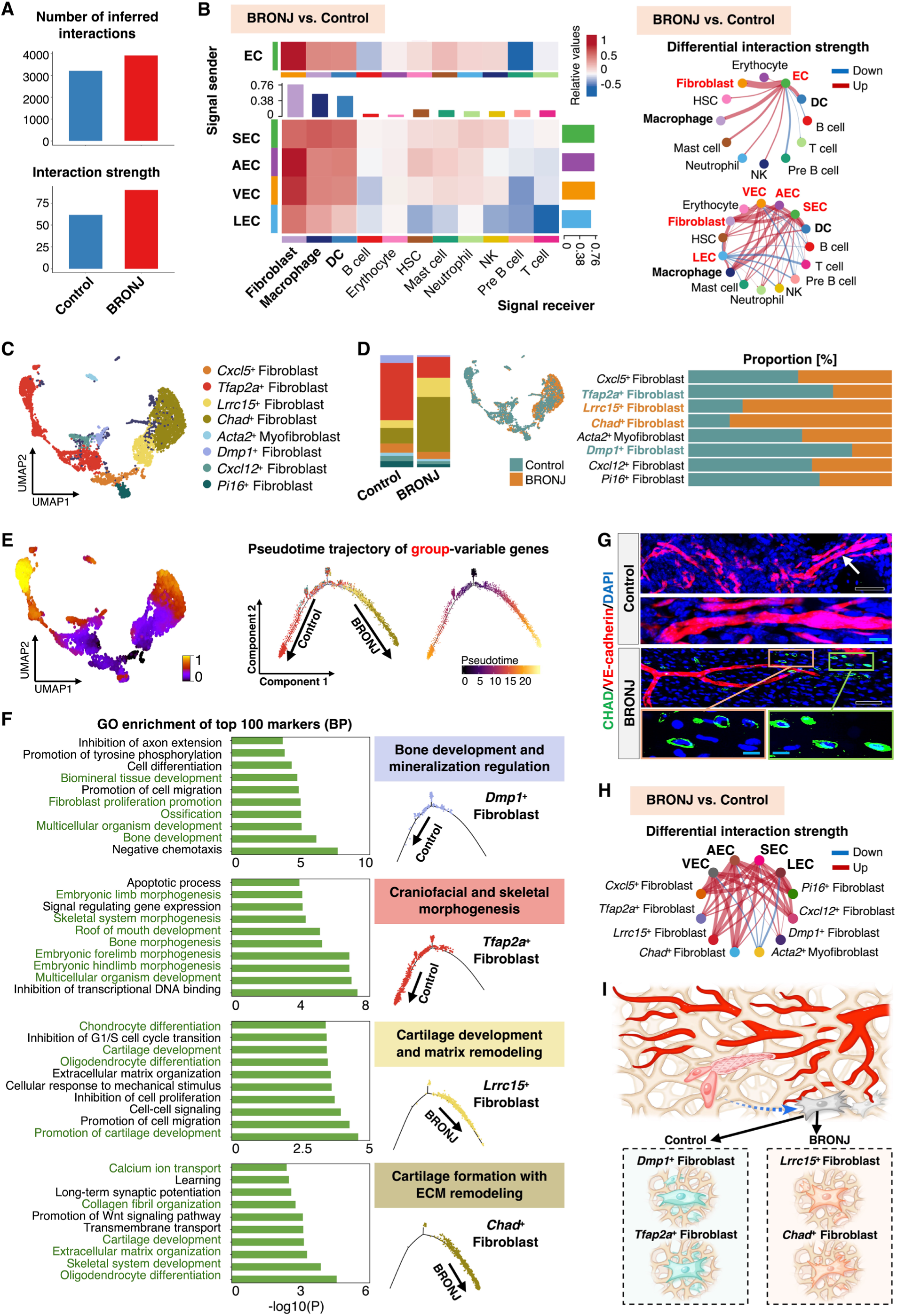
Fibroblast heterogeneity and endothelial-stromal dysregulation. (**A**) Changes in the number and strength of intercellular interactions between control and BRONJ groups by CellChat analysis. (**B**) Heatmap and chord diagram of communication changes, with EC subtypes as signal senders and other cell clusters as receivers. (**C**) UMAP of fibroblast subtypes. (**D**) Proportional changes of fibroblast subtypes in control and BRONJ groups. (**E**) RNA velocity by scVelo and pseudotime trajectories inferred by Monocle2. (**F**) GO enrichment of the top 100 marker genes for four differentially represented fibroblast subtypes. (**G**) Imaging of extraction sockets from control and BRONJ groups stained for CHAD and VE-cadherin, with nuclei counterstained with DAPI; arrows indicate magnified regions. (**H**) Chord diagrams of interaction strength between EC and fibroblast subtypes in control and BRONJ groups. (**I**) Schematic single-cell atlas of fibroblast subtypes in the endothelial microenvironment of control and BRONJ mandibles. **Scale bars**: black, 50 µm; and blue, 20 µm.

Similar to ECs, fibroblasts display pronounced heterogeneity (*29, 30*), encompassing phenotypic differences between healthy and diseased states as well as functional specialization shaped by the stromal microenvironment (*31*). To further investigate fibroblast pathology in the BRONJ microenvironment, we identified 8 subtypes, including *Cxcl5*⁺, *Tfap2a*⁺, *Lrrc15*⁺, *Chad*⁺, *Acta2*⁺, *Dmp1*⁺, *Cxcl12*⁺, and *Pi16*⁺ fibroblasts (Fig. 4C, and fig. S6C). In controls, *Tfap2a*⁺ and *Dmp1*⁺ fibroblasts predominated, whereas in BRONJ, *Lrrc15*⁺ and *Chad*⁺ fibroblasts selectively expanded (Fig. 4D). Despite a global reduction in intercellular communication, *Lrrc15*⁺ and *Chad*⁺ fibroblasts exhibited abnormally heightened signaling interactions (fig. S6D). Trajectory analysis indicated their progression toward a pathological terminal state characterized by aberrant chondrogenesis and matrix imbalance upon BRONJ condition (Fig. 4, E and F, and fig. S6, E and F), further corroborated by immunofluorescence detection of expanded CHAD⁺ fibroblasts in BRONJ tissues (Fig. 4G, and fig. S6G). Notably, this fibroblast subset has also been identified in previous studies and termed matrifibrocytes (*32*), characterized by the expression of cartilage-remodeling genes such as *Chad*, *Comp*, and *Cilp2*, consistent with the marker profile observed in our study (fig. S6C). In contrast, *Tfap2a*⁺ and *Dmp1*⁺ fibroblasts in controls were predominantly associated with bone development and morphogenesis (Fig. 4F). These findings reveal that fibroblasts undergo pathological chondrogenic remodeling, leading to aberrant cartilage-like matrix deposition and remodeling.

Moreover, CellChat analysis demonstrated significantly enhanced communication between most endothelial subtypes and fibroblast populations in BRONJ (Fig. 4H). Accordingly, we constructed a single-cell atlas of EC-fibroblast interactions within the bone microenvironment (Fig. 4I), which clearly delineates fibroblast remodeling toward a pathological chondrogenic state during BRONJ. Overall, these results indicate that BRONJ stems from a complex pathological network intertwining endothelial decline with stromal abnormalities, rather than from a single cell type dysfunction.

### EC-macrophage proinflammatory coupling and immune imbalance in BRONJ

The transcription factor (TF) network of ECs, as a central regulator of fate determination and functional plasticity, plays a pivotal role in maintaining vascular homeostasis and driving pathological transitions (*33*). SCENIC analysis revealed that ECs in controls were enriched for angiogenesis-related TF, whereas BRONJ ECs were markedly enriched for immune and inflammation-related TF (Fig. 5A), indicating a shift from a vascular homeostatic to a proinflammatory transcriptional program, accompanied by strengthened EC-macrophages/DCs communication in BRONJ (Fig. 5B). Concurrently, macrophage lineages exhibited widespread upregulation of inflammatory genes in BRONJ (fig. S7, A to C), closely mirroring the proinflammatory transcriptional profile of ECs (Fig. 5C). Together, these findings define an aberrantly amplified EC-macrophage proinflammatory coupling axis in BRONJ.

**Fig. 5.**
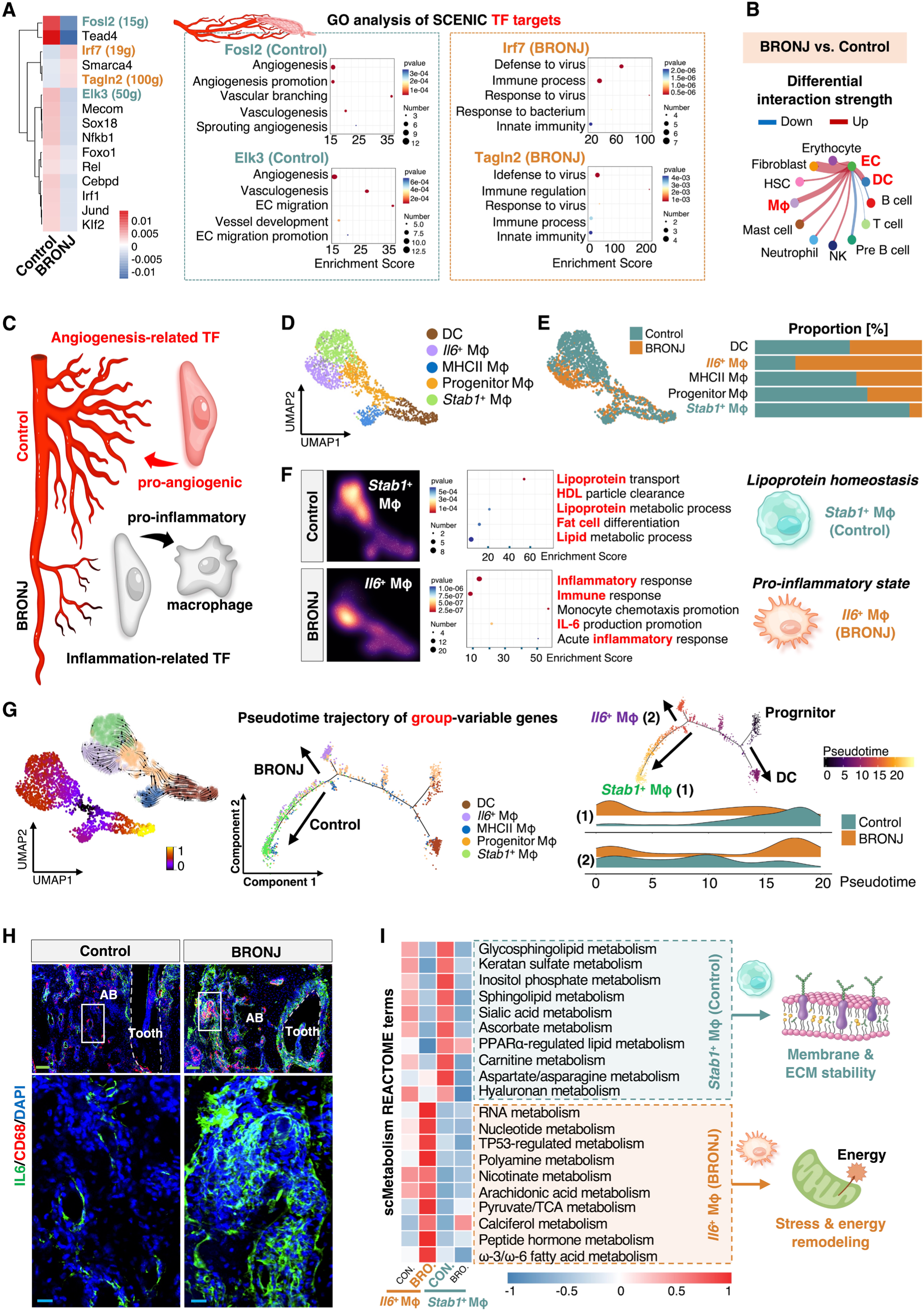
EC-macrophage proinflammatory remodeling in BRONJ. (**A**) SCENIC regulon analysis of ECs between groups and GO enrichment of TF target genes. (**B**) Chord diagram of ECs as signal senders and other clusters as receivers. (**C**) Schematic of EC transcription factors based on SCENIC and CellChat results. (**D**) UMAP of macrophage subtypes. (**E**) Proportional changes of macrophage subtypes in control and BRONJ groups. (**F**) GO enrichment of the top 100 marker genes for *Stab1*⁺ and *Il6*⁺ macrophages. (**G**) RNA velocity by scVelo and pseudotime trajectories by Monocle2. (**H**) Imaging of control and BRONJ extraction sockets stained for IL6 and CD68, with nuclear counterstaining by DAPI. (**I**) scMetabolism reactome heatmap and functional schematic of *Stab1*⁺ and *Il6*⁺ macrophages. **Scale bars:** green, 100 µm; and blue, 20 µm.

Subclustering defined 5 macrophage states: progenitors, DCs, MHCII⁺ macrophages, *Il6*⁺ macrophages, and *Stab1*⁺ macrophages (Fig. 5D, and fig. S7D). In healthy controls, *Stab1*⁺ macrophages predominated and were linked to lipoprotein homeostasis (Fig. 5, E and F), consistent with prior studies showing their involvement in lipid homeostasis and defense against oxidative tissue injury (*34*). In BRONJ this population was replaced by *Il6*⁺ macrophages, which expanded markedly and displayed a strongly inflammatory phenotype (Fig. 5, E and F). Pseudotime analysis of the macrophage lineage revealed that progenitors bifurcated into DCs and mature macrophages, with the latter skewing toward terminal *Stab1*⁺ states in controls but toward *Il6*⁺ states in BRONJ (Fig. 5G). This lineage framework alighs with recent single-cell studies showing that mononuclear phagocytes, including macrophages and DCs, arise from a common progenitor (*35*) and, under microenvironmental cues, differentiate into either homeostatic macrophages or inflammatory effector populations (*36*). Immunolabeling also validated elevated IL-6 in BRONJ lesions (Fig. 5H), indicating a proinflammatory microenvironment. Macrophages adapt their metabolism to homeostatic versus disease contexts (*37*), as reflected by our metabolic profiling results (Fig. 5I). Control *Stab1*⁺ macrophages maintained membrane and ECM homeostasis via sphingolipid and polysaccharide metabolism, whereas BRONJ *Il6*⁺ macrophages activated energy metabolism, indicating stress-driven reprogramming toward a hyperinflammatory state (Fig. 5I).

To further delineate immune dysregulation, we identified 8 NK/T cell subtypes. All were markedly reduced in BRONJ except *Cd8*⁺ T cell (fig. S7, F and G). In parallel, B cells and pre-B cell populations also declined (Fig. 2C), indicating disruption of both innate and adaptive compartments (*38*). Thus, immune imbalance in BRONJ manifests as a multidimensional process: ECs reprogrammed from angiogenic to inflammatory states; macrophages shifted toward proinflammatory fates with stress-responsive metabolic remodeling; and immune regulatory networks were broadly compromised.

### *Col2a1* signaling, rather than EndoMT, mediates EC-driven stromal and immune dysregulation in BRONJ

Having delineated endothelial attrition and fibroblast/macrophage shifts in BRONJ, we next focused on cross-cellular signaling. CellChat analysis revealed globally reinforced communication from all 4 EC subtypes to fibroblast/macrophage states in BRONJ (Fig. 6A), with collagen signaling emerging as dominant (Fig. 6B). Notably, type I and II collagen signals were markedly enhanced, whereas type IV collagen signaling was diminished (fig. S8A), with histology confirming Collagen IV loss in BRONJ basement membranes (fig. S8, B and C). Among the upregulated axes, the COL2A1-CD44 axis (EC-to-fibroblast/macrophage signaling) and COL2A1-SDC4 (EC-to-fibroblast signaling) axis was uniquely BRONJ-specific (Fig. 6C, and fig. S8D). *Col2a1* was markedly elevated across all EC subtypes, exceeding the induction of other genes (Fig. 6D), while perivascular Collagen II deposition accumulated within the BRONJ extraction sockets (Fig. 6E). Immunolabeling at single-cell resolution revealed rare colocalization of Collagen II with ECs (fig. S8E). Reduced Collagen IV reflected loss of endothelial features, whereas abnormal Collagen I/II elevation suggested fibroblast-like secretory activity, which prompted consideration of endothelial-to-mesenchymal transition (EndoMT) (*39*). While EndoMT is essential in embryonic heart development (*40*) and has been implicated in pulmonary fibrosis (*41*) and cerebrovascular inflammation (*42*), recent evidence suggests protential roles in marrow homeostasis (*43*), yet its contribution to physiological and pathological processes in bone remains unresolved (*44*). We therefore investigated whether vascular-stromal remodeling in BRONJ is EndoMT-dependent or instead driven by alternative mechanisms.

**Fig. 6.**
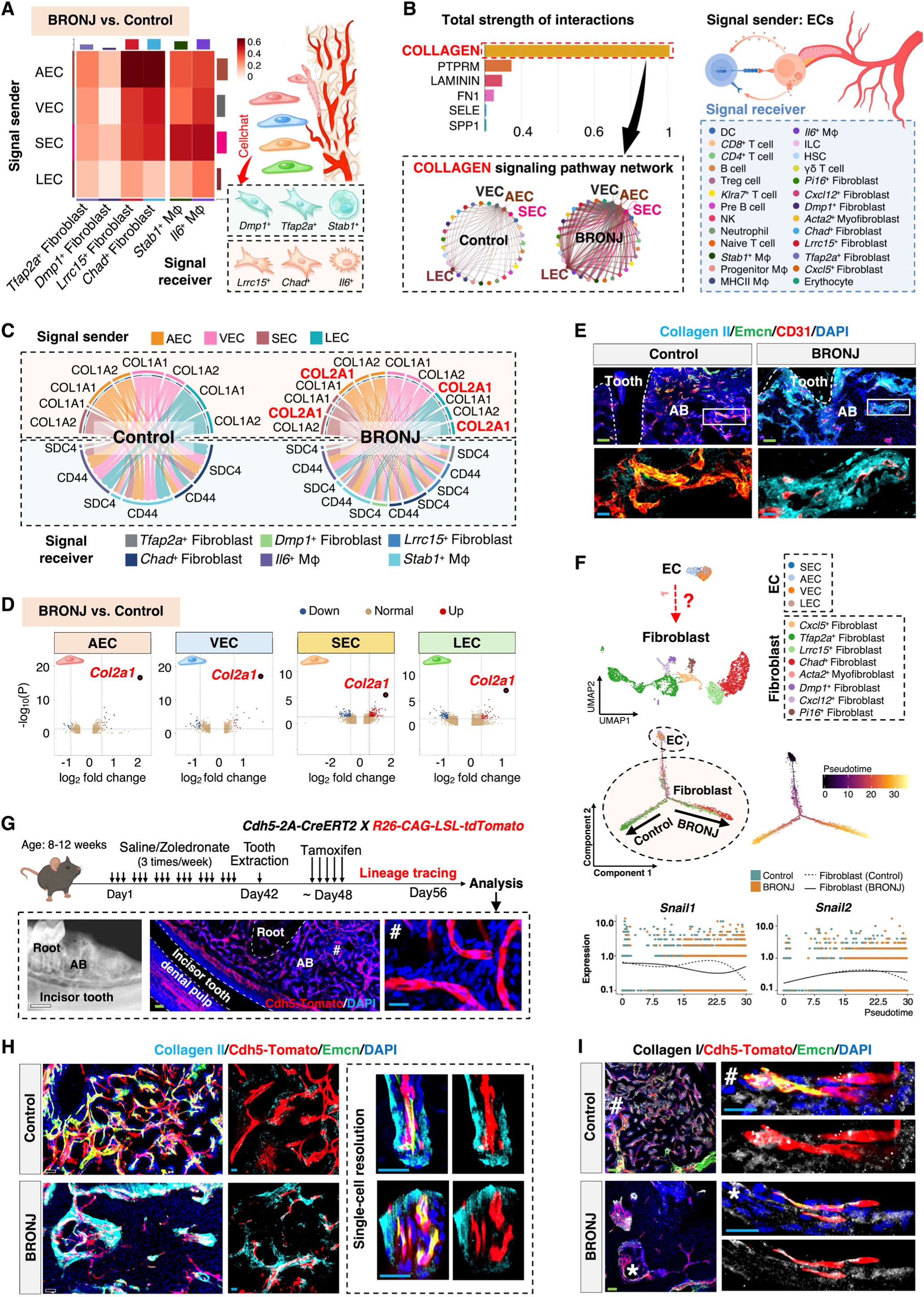
Endothelial collagen signaling and EndoMT lineage tracing in BRONJ. (**A**) Altered signaling from EC subtypes to key fibroblast and macrophage populations. (**B**) Pathway changes in BRONJ highlighting the collagen network. (**C**) Ligand-receptor pairs of collagen I/II pathways among major cell subtypes. (**D**) Volcano plots of differentially expressed genes in the four EC subtypes. (**E**) Collagen II expression in the extraction socket with vessels marked by CD31 and Emcn. (**F**) Monocle2 trajectory of the integrated EC-fibroblast UMAP evaluating EndoMT transcription factors *Snail1* and *Snail2*. (**G**) Lineage tracing in Cdh5-2A-CreERT2 × R26-CAG-LSL-tdTomato BRONJ mice, showing mandibular structures by X-ray and Cdh5-tdTomato thick sections with DAPI nuclear counterstaining. (**H**) High-resolution imaging of endogenous Cdh5⁺ vessels in extraction sockets of control and BRONJ mice stained with Collagen II and Emcn, with nuclei counterstained by DAPI. (**I**) Imaging of Cdh5⁺ vessels stained with Collagen I and Emcn, with nuclei counterstained by DAPI. **Scale bars:** white, 500 µm; green, 100 µm; black, 50 µm; and blue, 20 µm.

Based on pseudotime trajectories of the EC-fibroblast continuum, no significant induction of canonical EndoMT markers such as *Snail1* and *Snail* 2 (Fig. 6F) or EndoMT marker-based scoring (fig. S9A) was observed in the inferred transition zone. In parallel, however, re-clustering of the EC-fibroblast continuum identified a mixed population enriched for angiogenic and stromal remodeling signatures, resembling an EndoMT-like intermediate state (fig. S9B). To test this possibility, we performed lineage tracing in *Cdh5-2A-CreERT2*; *Rosa26-LSL-td-Tomato* BRONJ mice (Fig. 6G), and immunolabeling with colocalization analysis revealed that over 99% of Collagen I and II deposits were tdTomato⁻ in BRONJ (Fig. 6, H and I, and fig. S9C), providing strong evidence against EndoMT. Thus, in BRONJ, ECs drive pathology not through EndoMT but via aberrant *Col2a1* signaling, acting as a paracrine hub through the COL2A1-CD44/SDC4 axis to promote ECM remodeling and inflammation.

### Cross-species validation of vascular-lymphatic abnormalities in BRONJ

In the murine model, we identified a pathological axis characterized by endothelial loss and stromal imbalance. To validate and extend these findings, we performed integrated proteomic profiling and vascular-lymphatic imaging in human BRONJ samples (Fig. 7A). Proteomic analysis revealed a systemic attenuation pattern, with markedly reduced total protein numbers, widespread downregulation, and limited upregulation (fig. S10A). Downregulated proteins were enriched in angiogenic and osteogenic functions, whereas upregulated proteins clustered in inflammatory pathways (Fig. 7B), revealing enhanced inflammation, endothelial dysfunction, and impaired bone formation. Notably, matrix metalloproteinases (MMPs) emerged as the upregulated hub in the protein-protein interaction network, potentially driving extracellular matrix remodeling and disrupting endothelial-immune homeostasis by facilitating leukocyte transendothelial migration (fig. S10B). The MMP family degrades basement membrane components (*45*) such as proteoglycans (*46*) and extracellular matrix fibrils including type II collagen (*47*), ultimately inducing structural disorganization and pathological remodeling upon BRONJ.

**Fig. 7.**
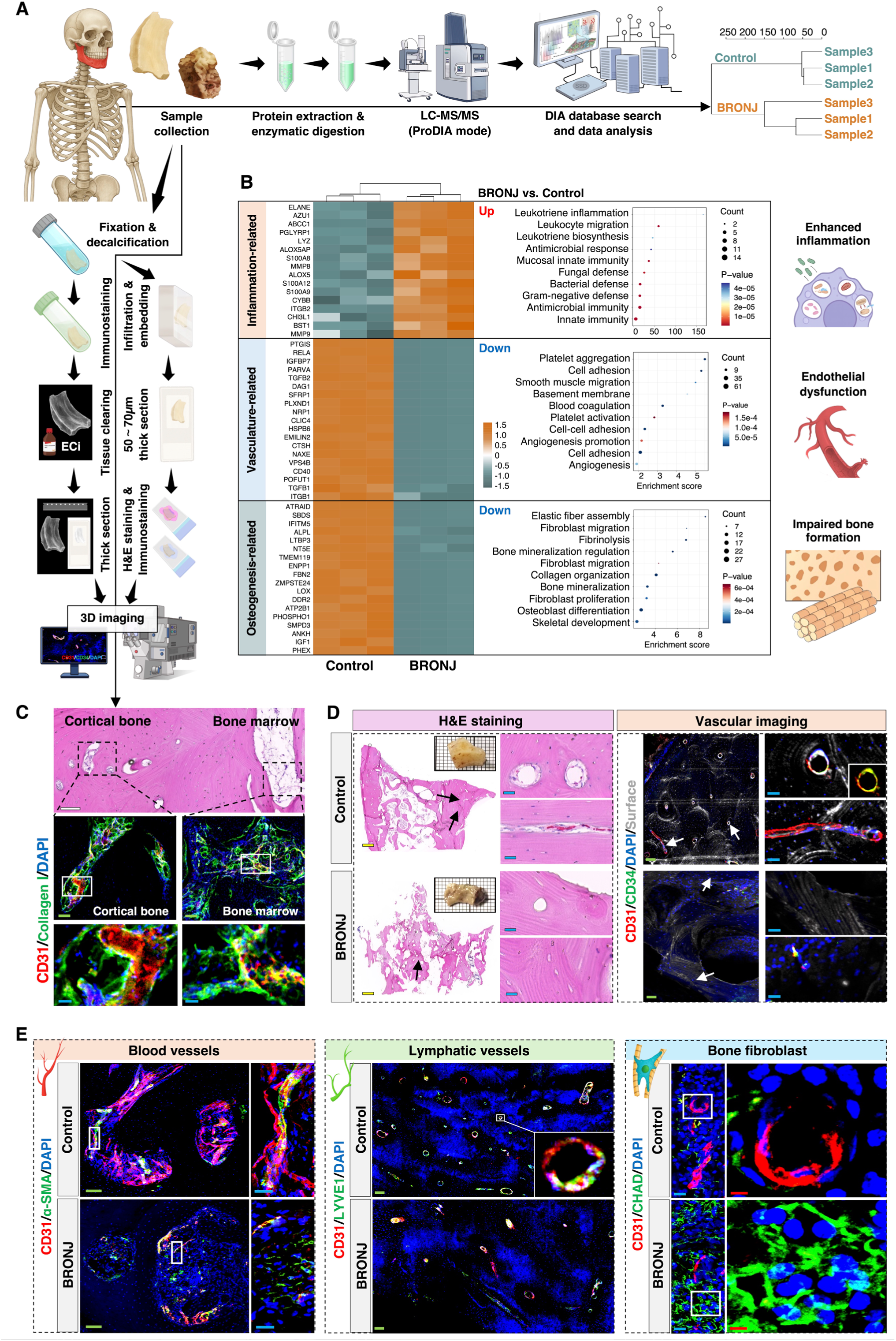
Proteomic and histological characterization of human BRONJ samples. (**A**) Workflow of proteomic analysis and histology/immunofluorescence in human mandibular specimens *(n = 3)*. (**B**) Heatmap of differentially expressed inflammation-, vasculature-, and osteogenesis-related proteins, with GO enrichment of upregulated and downregulated proteins in BRONJ. (**C**) Histological and fluorescence imaging of cortical bone and marrow; stained for CD31 and Collagen I, with nuclei counterstained by DAPI. (**D**) Histological and fluorescence imaging of control and BRONJ samples; stained with CD31 and CD34, with nuclei counterstained by DAPI; arrows indicate magnified regions. (**E**) Images of control and BRONJ samples, with blood vessels stained for CD31 and α-SMA, lymphatic vessels for CD31 and LYVE1, fibroblasts for CD31 and CHAD, and nuclei counterstained with DAPI. **Scale bars:** yellow, 1000 µm; white, 500 µm; green, 100 µm; blue, 20 µm; red, 5 µm. n represents biological replicates.

Although anti-angiogenic effects of bisphosphonates are widely postulated to underlie human BRONJ, current evidence largely derives from animal studies (*48*) or indirect mucosal microvascular imaging at the margins of osteonecrotic lesions (*49*), leaving “bone angiogenesis inhibition” a consensus hypothesis lacking direct visualization. Compared with mice, human cortical bone is denser, with a more complex marrow architecture and interspecies differences in endothelial and stromal markers, enhancing imaging difficulty. By overcoming antigen loss from conventional acid decalcification and integrating tissue-clearing steps, we established a 3D imaging method for the vascular network in the human bone (Fig. 7, A and C, and fig. S10C). Histology confirmed typical BRONJ osteonecrotic lesions and revealed endothelial injury (Fig. 7D). For the vascular compartment, CD31⁺CD34⁺ microvessel density was markedly reduced (Fig. 7D), and CD31⁺α-SMA⁺ arterial vessels were compromised (Fig. 7E, and fig. S10D). Regarding the lymphatic system, CD31⁺LYVE1⁺ lymphatic vessels also exhibited a marked reduction in human BRONJ bone (Fig. 7E). Moreover, pathological expansion of CHAD⁺ fibroblasts was again observed in BRONJ (Fig. 7E), further validating the endothelial-stromal disorder in a cross-species context.

In this study, by integrating high-resolution 3D imaging, single-cell transcriptomics, and proteomics, we comprehensively profiled murine models and human mandibular specimens, generating a cross-species atlas of the jawbone microenvironment (Fig. 8). At the tissue level, our findings demonstrate gross vascular and lymphatic injury in BRONJ. At the single-cell scale, 4 endothelial, 4 fibroblast, and 2 macrophage subtypes were identified as key drivers to BRONJ progression; and at the molecular level, COL2A1-CD44 axis (EC-to-fibroblast/macrophage signaling) and COL2A1-SDC4 (EC-to-fibroblast signaling) axis orchestrated the vascular, stromal, and immune pathological networks (Fig. 8). This integrated framework not only refines our mechanistic understanding but also highlights potential avenues for therapeutic intervention.

**Fig. 8.**
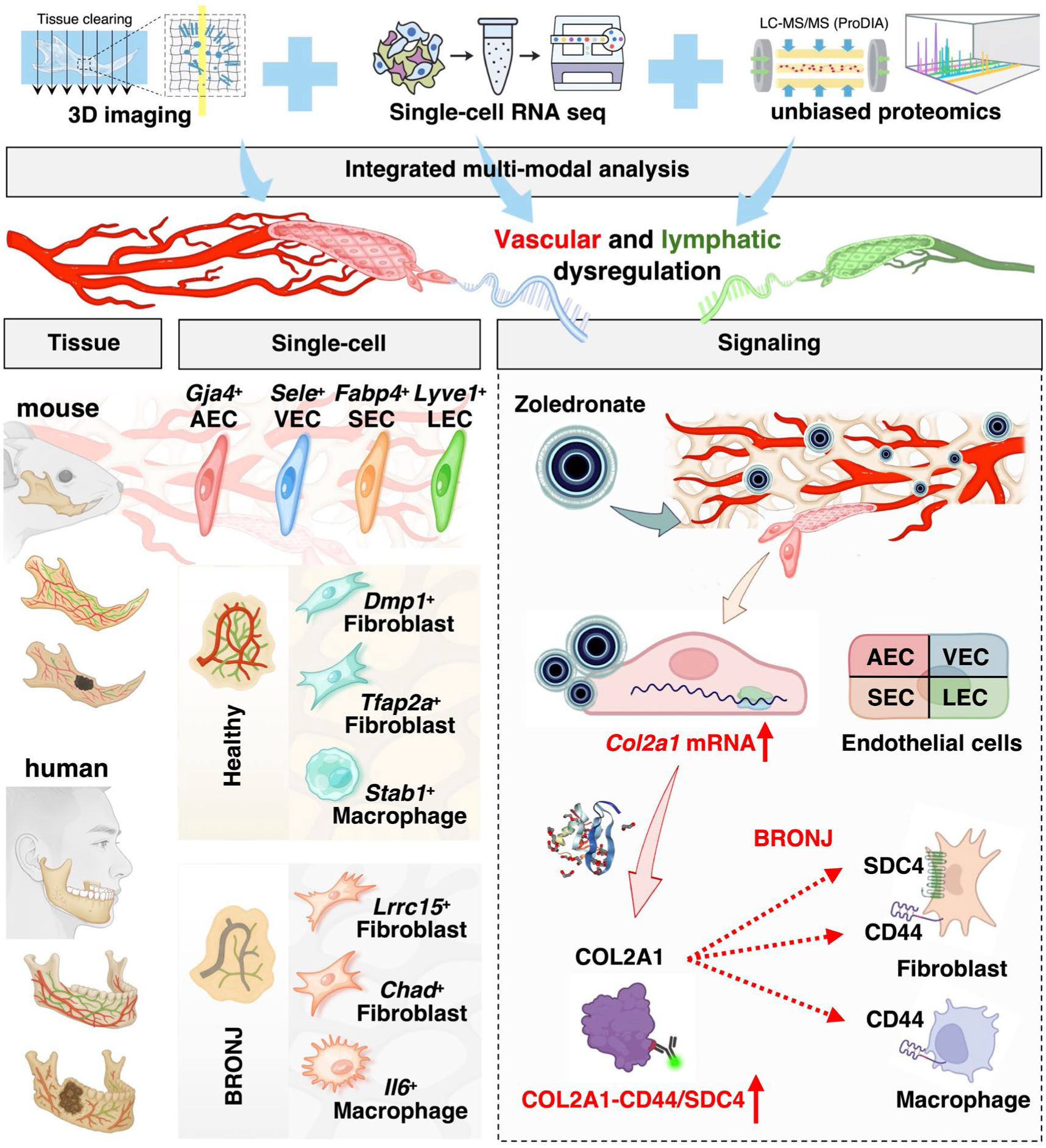
Schematic of the vascular-lymphatic dysregulated microenvironment in BRONJ. Integrating high-resolution tissue-clearing imaging, single-cell transcriptomics, and proteomics, the schematic depicts vascular-lymphatic dysregulation in BRONJ, with EC subtype shifts, stromal-immune remodeling, and zoledronate-induced COL2A1-CD44/SDC4 axis.

## Discussion

BRONJ represents one of the most severe complications of antiresorptive therapy, with a substantial clinical burden that is well recognized. However, the proposed mechanism of “impaired angiogenesis” has so far rested on fragmented evidence (*6–8, 11, 14, 15*), lacking systematic validation or a coherent framework. Our findings demonstrate that vascular and lymphatic endothelial injury does not occur in isolation but is tightly interwoven with aberrant matrix remodeling and immune-inflammatory imbalance, together establishing a central pathological axis that drives BRONJ progression.

Through whole tooth-jawbone clearing and 3D imaging, this study uncovered profound vascular and lymphatic rarefaction in BRONJ. Single-cell transcriptomics further demonstrated transcriptional reprogramming across all endothelial subtypes, most notably the global upregulation of *Col2a1*, implicating ECs not only as direct targets of zoledronate toxicity but also as active drivers of pathological matrix remodeling. We delineated an integrated pathological network characterized by vasculature collapse, aberrant chondrogenic fibroblasts and *Il6*⁺ macrophage infiltration. Notably, IL6 enrichment in BRONJ sockets echoes its role in periodontitis as a key mediator of bone destruction (*50*), suggesting IL6 as an amplifier of “matrix-immune” injury. Moreover, although increased Collagen I/II deposition alongside reduced Collagen IV suggested possible EndoMT, lineage tracing definitively excluded this, demonstrating that a non-EndoMT paracrine program represents the operative mechanism.

This study also has certain limitations. First, BRONJ is a progressive disease, and although early endothelial loss and stromal abnormalities were evident within 2 weeks post-extraction, the initial vascular remodeling events remain to be elucidated. In addition, systemic metabolic and endocrine factors warrant consideration (*51*). Early postmenopausal estrogen decline not only regulates type H vessel physiology (*52*), but also interacts with zoledronate in the immune microenvironment (*53*), potentially shaping both drug efficacy and tissue response. Thus, investigations of BRONJ would benefit from considering bisphosphonate pharmacokinetics and early postmenopausal hormonal effects on bone vasculature and immunity. Moreover, using human samples, this study confirmed strong concordance with murine findings while revealing certain human-specific features, highlighting the need for larger cohorts to further elucidate BRONJ mechanisms and therapeutic targets.

From a translational perspective, our findings highlight three complementary BRONJ therapeutic avenues. First, vascular protection and basement membrane reconstruction, including promotion of angiogenesis, biomaterial-based scaffolds, or Collagen IV preservation. Second, rebalancing the stromal-immune microenvironment by reversing chondrogenic fibroblast phenotypes and proinflammatory macrophage states. Moreover, blockade of endothelial paracrine signaling, particularly via interruption of the COL2A1-CD44/SDC4 axis.

Overall, our study reframes BRONJ from a simplistic view of angiogenic inhibition to a multidimensional disease characterized by vascular-lymphatic regression, matrix remodeling, and immune imbalance. Moreover, we identify the COL2A1-CD44/SDC4 axis as a central therapeutic target that links the vascular, stromal, and immune compartments. Interventions aimed at endothelial repair and paracrine signaling holds promise to shift BRONJ management from passive debridement toward active restoration of microenvironmental homeostasis.

## Methods

### Mouse models

Female C57BL/6J mice (8-12 weeks old) were purchased from Dashuo Biotechnology Co., Ltd. (Chengdu, China). To establish a murine model of bisphosphonate-related osteonecrosis of the jaw (BRONJ), zoledronate (200 μg/kg; MCE, HY-13777) or an equal volume of saline (vehicle control) was administered intraperitoneally three times per week from day 1 to day 56. Extraction of bilateral 1^st^ mandibular molars was performed on day 42. Mice were euthanized on day 56, and mandibles were harvested.

*Cdh5-2A-CreERT2* (stock no.NM-KI-200173) and *Rosa26-LSL-td-Tomato* (stock no.NM-KI 225042) mice were obtained from Shanghai Southern Model Biotechnology Co., Ltd. (China). To achieve genetic labeling, *Cdh5-2A-CreERT2* transgenic mice were crossed with *Rosa26-LSL-td-Tomato* reporter mice. For tamoxifen induction, tamoxifen (Sigma-Aldrich, T5648) was dissolved in corn oil at 20 mg/ml. *Cdh5-2A-CreERT2*; *Rosa26-LSL-tdTomato* transgenic mice received oral tamoxifen at 75 mg/kg daily for 5 consecutive days to activate Cre recombinase. Mice were sacrificed 9 days after the final administration. All animal experiments complied with local ethical guidelines and regulations and received approval from the Research Ethics Committee of West China Hospital of Stomatology (Approval No. WCHSIRB-D-2023-245).

### Human samples

Mandibular bone specimens were collected from patients undergoing segmental mandibulectomy at West China Hospital of Stomatology (Chengdu, China). All experiments using these human samples received approval from the Research Ethics Committee of West China Hospital of Stomatology (Approval No. WCHSIRB-D-2025-392). Written informed consent was obtained from all participants.

### micro-CT

Mandibles were dissected, stripped of soft tissues, and fixed in 4% paraformaldehyde (PFA; Sigma-Aldrich, P6148). High-resolution micro-CT scans were performed using a SCANCO μCT50/100 system with the following parameters: voxel size, 10.0 μm; field of view, 34 mm × 110 mm; scan length, 6.220 mm (83.857-90.077 mm); voltage, 70 kVp; current, 200 μA; integration time, 300 ms.

### Histology (H&E staining)

Samples were dehydrated through graded ethanol, cleared in xylene, and embedded in paraffin. Sections (5 μm) were cut, deparaffinized, rehydrated, and stained with hematoxylin (5 min), differentiated in acid alcohol, blued in running tap water, and counterstained with eosin (2 min). Images were acquired with a digital slide scanner (Olympus SLIDEVIEW VS200).

### Tissue processing, fixation, and decalcification

Freshly dissected mandibles were fixed in 4% PFA on ice for 2.5 h after removal of muscle, periosteum, and other soft tissues. Decalcification was performed with 0.5 M EDTA at room temperature for 2 days with buffer refreshed twice daily, followed by three PBS washes.

### Whole-organ tissue clearing and immunostaining of jawbone

Decalcified mandibles were dehydrated in graded cold methanol (50% and 80% for 30 min each, followed by 100% for 1 h with two changes every 20 min). Tissues were bleached in 5% H₂O₂ (Sigma-Aldrich, HX0636) for 1 h, rehydrated through a descending methanol series, and washed in PBS. Permeabilization was performed in a buffer containing 25% urea (Sigma-Aldrich, U5128), 15% glycerol (Sigma-Aldrich, G7893), and 15% Triton X-100 (Sigma-Aldrich, T8787) at 4 °C for 5 h, followed by digestion with 0.2% collagenase (Merck, 10103578001) at 37 °C for 30 min. After blocking with 10% donkey serum, 10% DMSO, and 0.5% Triton X-100 in PBS at 37 °C for 20 min, samples were incubated overnight with primary antibodies diluted in 2% donkey serum/10% DMSO/0.5% Triton X-100 in PBS. After extensive washes, fluorophore-conjugated secondary antibodies were applied at 37 °C for 6-8 h, followed by additional washes. Samples were dehydrated through graded isopropanol (30%, 50%, 80%, 100%), cleared in ethyl cinnamate (ECi; Sigma-Aldrich, 112372), and further equilibrated in 80% ECi/20% PEGM (Sigma-Aldrich, 447943). Imaging was performed on a Zeiss Lightsheet 7 microscope with a 5× EC Plan-Neofluar objective. Images were processed using Imaris File Converter and Imaris (v10.2.0). All antibody information is listed in Supplementary Table 1.

### Thick-section immunofluorescence imaging

Decalcified bone tissues were incubated in 20% sucrose (Sigma-Aldrich, V900116) and 2% polyvinylpyrrolidone (PVP; Sigma-Aldrich, P5288) at 4 °C for 12 h, followed by embedding in 8% gelatin (Sigma-Aldrich, G1890) supplemented with 20% sucrose and 2% PVP. Samples were sectioned with 70∼100 μm thickness using a Leica CM1950 cryostat (Leica Microsystems, Germany). After rehydration, sections were permeabilized with 0.3% Triton X-100 and blocked with 5% donkey serum. Primary antibodies were applied at a 1:150 dilution and incubated for 4 h at room temperature, followed by Alexa Fluor-conjugated secondary antibodies (1:300) for 1-1.5 h. Nuclei were counterstained with DAPI (Solarbio, C0060), and sections were mounted in Fluoromount-G (Invitrogen, 00-4958-02) and stored at 4 °C until imaging. Images were acquired on a spinning-disk confocal microscope (Olympus, IXplore IX83 SpinSR) equipped with 405, 488, 561, and 647 nm laser lines. Imaging modalities included multichannel acquisition, panoramic stitching, extended focus, z-stacks, and time-lapse imaging. Image processing and three-dimensional reconstruction were performed with Imaris software (v10.2.0).

### Single-cell RNA sequencing (scRNA-seq)

Freshly dissected mandibular bones were rinsed in PBS, minced (∼0.5 mm³), and digested with 0.2% collagenase I (Gibco, 17018029), II (Gibco, 17101015), and IV (Gibco, 17104019) for 40 min, followed by 0.25% trypsin (Gibco, 15090046) for 10 min at 37 °C. Cell suspensions were filtered through a 40 μm strainer (BD), centrifuged (300 g, 5 min, 4 °C), and subjected to red blood cell lysis (MACS, 130-094-183). CD45⁻ cells were enriched using magnetic beads (MACS, 130-052-301) (*54, 55*). Viable cells were counted (Luna cell counter), resuspended at 700-1200 cells/μL, and loaded into the 10× Genomics Chromium platform using the Next GEM Single Cell 3′ v3.1 kit (1000268). Libraries were sequenced on an Illumina NovaSeq 6000 PE150 system (OE Biotech, Shanghai). Raw sequencing data were processed with Cell Ranger (*56*) (v7.0.1) for alignment to the mm10 genome, barcode filtering, and UMI quantification. Downstream analysis was performed in Seurat (v4.0.0). high-quality cells were defined as those with gene and UMI counts within the median ± 2 median absolute deviations (MADs) and with mitochondrial UMI fraction <20%. Doublets were subsequently removed using DoubletFinder (*57*) prior to downstream analysis. Normalization was performed with NormalizeData, highly variable genes (top 2000) were selected with FindVariableGenes, and dimensionality reduction was conducted by UMAP. Differentially expressed genes (DEGs) were identified using FindMarkers (test.use = presto), using thresholds of p < 0.05 and fold change >1.5. GO and KEGG enrichment analyses were conducted by hypergeometric testing.

### Cell-cell communication analysis

Cell-cell ligand-receptor interactions were analyzed with the Cellchat (*58*) R package (version 1.1.3). The normalized expression matrix was first imported, and a Cellchat object was constructed using the createCellchat function. Preprocessing was performed with default parameters using the functions identifyOverExpressedGenes, identifyOverExpressedInteractions, and projectData. Potential ligand-receptor interactions were then inferred with computeCommunProb, filtered using filterCommunication (min.cells=10), and further analyzed with computeCommunProbPathway. Finally, intercellular communication networks were aggregated via the aggregateNet function.

### Pseudotime trajectory analysis with Monocle2

Cell differentiation trajectories were reconstructed using the Monocle2 (*59*) R package (version 2.9.0). Briefly, Seurat objects were first converted into CellDataSet objects with the importCDS function. Ordering genes were identified using the differentialGeneTest function (q-value<0.01). Dimensionality reduction and clustering were then conducted with the reduceDimension function, followed by trajectory inference using the orderCells function.

### RNA velocity analysis with scVelo

Based on the Cell Ranger output, spliced and unspliced reads were quantified using the Python package velocyto.py (*60*). RNA velocity was then estimated with scVelo (*61*) by constructing a likelihood-based dynamical model to infer gene-specific transcription, splicing, and degradation rates. This approach enabled the reconstruction of each cell’s position along potential differentiation trajectories and the calculation of RNA velocities at the single-cell level, which were subsequently projected onto t-SNE or UMAP embeddings for visualization.

### SCENIC analysis

Gene regulatory network inferences were performed through SCENIC (v1.2.4) (*62*), RcisTarget (v1.10.0), and AUCell (v1.12.0). Co-expression modules were identified, motifs were validated with RcisTarget, and regulon activity was scored with AUCell. Regulon specificity scores (RSS) (*63*) were computed with scFunctions (https://github.com/FloWuenne/scFunctions/) based on Jensen-Shannon divergence.

### Metabolic activity analysis

Metabolic pathway activity at single-cell resolution was quantified with scMetabolism (*64*) (v0.2.1), using KEGG (85 pathways) and Reactome (82 pathways) gene sets. Homology conversion was applied to mouse data, and activity scores were computed using the VISION (*65*) algorithm.

### Module score analysis with AddModuleScore

Module scores for endothelial, EndoMT, and fibroblast gene sets were calculated using the AddModuleScore function in the Seurat R package. The function computes the average expression of genes within each set at the single-cell level and corrects it by subtracting the mean expression of matched control genes randomly sampled from bins of genes with comparable expression levels.

### LC-MS/MS high-resolution proteomics

Frozen jawbone tissues were ground in liquid nitrogen, and proteins were extracted using the phenol-methanol method with phosphatase and protease inhibitors. After precipitation/washing with ammonium acetate-methanol and acetone, protein pellets were dissolved, quantified by BCA, and quality-checked by SDS-PAGE. Equal amounts were captured with SP3 magnetic beads, reduced with DTT, alkylated with CAA, and digested with trypsin at 37 °C. Peptides were desalted, spiked with iRT standards, and analyzed by LC-MS/MS in DIA mode. Separation was performed on C18 columns with a linear gradient of 0.1% formic acid in water and acetonitrile. High-resolution spectra were acquired on an Orbitrap platform and processed using DIA-NN. Proteins were retained if supported by ≥1 unique peptide and ≥2 valid values in ≥50% of samples in at least one group. Missing values were imputed by within-group means or half-minimum replacement, followed by median normalization and log₂ transformation to generate reliable protein matrices for downstream statistical and pathway analyses.

### Image analysis

Vessel density, integrated density, and signal colocalization were analyzed using Imaris (version 10.2) and Fiji (version 2.14.0/1.54f). For vessel density and integrated density, thresholds were adjusted in Fiji to define both the signal-positive area and the entire bone ROI. Signal quantification was calculated as the ratio of the signal-positive area to the total ROI area. 3D colocalization was assessed using the Imaris Coloc function. Briefly, a new surface was generated for one selected channel, and masking was applied to set voxels outside the surface to 0 and those inside to 100, thereby generating a new masked channel. For colocalization analysis, two channels of interest were selected together with the masked channel as the ROI (threshold set at 50). The thresholds of the two selected channels were manually adjusted until all vessel-associated signals were captured. The percentage of colocalization was then reported in the Coloc Estimated Volume Statistics output.

### Statistics analysis

Statistical comparisons between the healthy and BRONJ groups were conducted using unpaired two-tailed Student’s t-tests. Additionally, comparisons between different areas in one sample were analyzed using paired two-tailed Student’s t-tests. Both were analyzed by GraphPad Prism (GraphPad Software, USA, version 9.0). for these analyses. All that Valueswere presented as violin plot. ^ns^P >0.05; *P < 0.05; **P < 0.01; and ***P < 0.001. n represents biological replicates.

## Data availability

Source data are provided in this paper. The remaining data are available within the Article, Supplementary Information, and Source Data file.

## Conflict of interests

The authors do not declare competing financial interests.

## Acknowledgments

We thank OE Biotech Co., Ltd (Shanghai, China) for providing Sc RNA-seq, Chunyan Guan and Kunyue Meng for assistance with bioinformatics analysis. This study was supported by the National Nature Science Foundations of China (Nos. 82270961, 82422021), Sichuan Science and Technology Program (No. 2023JDRC0018), and Young Elite Scientist Sponsorship Program by CAST (No. 2023QNRC001) to J.C.; European Research Council (StG: metaNiche, 805201), Ministry of Education (MOE) Singapore: MOE-SUG and MOE Academic Research Funds Tier 1 (#024983-00001) to A.P.K..

## Contributions

A.P.K. and J.C. performed funding and supervised the study. Y.S., H.S., and J.C. contributed to design, data acquisition, interpretation, performed all statistical analyses and drafted. Z.G., X.L., X.P., and J.W. contributed to data acquisition and drew schematic diagrams, A.P.K., J.C., F.Y., X.P., and Q.Y. critically revised the manuscript. All authors have read and agreed to the published version of the manuscript.

## Notes

### Competing Interest Statement

The authors have declared no competing interest.

